# Preclinical validation and kinetic modelling of the SV2A PET ligand [^18^F]UCB-J in mice

**DOI:** 10.1101/2024.07.10.602899

**Authors:** Liesbeth Everix, Filipe Elvas, Alan Miranda Menchaca, Vinod Khetarpal, Longbin Liu, Jonathan Bard, Steven Staelens, Daniele Bertoglio

**Affiliations:** Molecular Imaging Center Antwerp (MICA), University of Antwerp, Wilrijk, Belgium; Molecular Imaging and Radiology (MIRA), Wilrijk, Belgium; CHDI Management, Inc., the company that manages the scientific activities of CHDI Foundation, Inc. Princeton, NJ, USA; Bio-Imaging Lab, University of Antwerp, Wilrijk, Belgium; µNeuro Center for Excellence, University of Antwerp, Antwerp, Belgium

**Author notes:** **Corresponding author:** Prof. Daniele Bertoglio, Bio-Imaging Lab, University of Antwerp, Universiteitsplein 1, Wilrijk, Belgium.

**Keywords:** mouse model, positron emission tomography, synapse, synaptic density, synaptic vesicle protein 2A

## Abstract

Synaptic vesicle protein 2A (SV2A) is ubiquitously expressed in presynaptic terminals where it functions as a neurotransmission regulator protein. Synaptopathy has been reported during healthy ageing and in a variety of neurodegenerative diseases. Positron emission tomography (PET) imaging of SV2A can be used to evaluate synaptic density. The PET ligand [^11^C]UCB-J has high binding affinity and selectivity for SV2A but has a short physical half-life due to the ^11^C isotope. Here we report the characterization and validation of its ^18^F-labeled equivalent, [^18^F]UCB-J, in terms of specificity, reproducibility and stability in C57BL/6J mice. Plasma analysis revealed at least one polar radiometabolite. Kinetic modelling was performed using a population-based metabolite corrected image-derived input function (IDIF). [^18^F]UCB-J showed relatively fast kinetics and a reliable measure of the IDIF-based volume of distribution (*V*_T(IDIF)_). [^18^F]UCB-J specificity for SV2A was confirmed through a levetiracetam blocking assay (50 to 200 mg/kg). Reproducibility of the *V*_T(IDIF)_ was determined through test-retest analysis, revealing significant correlation (r^2^=0.773, *p*<0.0001). Time-stability analyses indicate a scan duration of 60 min to be sufficient to obtain a reliable *V*_T(IDIF)_. In conclusion, [^18^F]UCB-J is a selective SV2A ligand with optimal kinetics in mice. Further investigation is warranted for (pre)clinical applicability of [^18^F]UCB-J in synaptopathies.

## INTRODUCTION

Synaptic vesicle glycoproteins 2 (SV2) are a family of highly conserved proteins involved in vesicular dynamics. To date, three isoforms have been identified: SV2A, SV2B and SV2C^1^. The predominant isoform, SV2A, is ubiquitously present in all human brain regions with highest expression observed in the thalamus and basal ganglia^1^. Due to its expression pattern and major involvement in neurotransmitter release from presynaptic terminals, SV2A is a promising marker to evaluate synaptic density^2^.

Impairment of synaptic density, synaptopathy, has been reported over the course of healthy ageing, as well as in multiple neurodegenerative and neuropsychiatric diseases such as Alzheimer’s disease (AD), Parkinson’s disease (PD), schizophrenia, and Huntington’s disease (HD)^3, 4^. Assessing synaptic density through SV2A detection can be performed using positron emission tomography (PET), a non-invasive imaging technique employing radioligands directed towards a target of choice. To date, several ligands for SV2A have been developed, including [^11^C]UCB-J, [^18^F]UCB-H, [^11^C]UCB-A, [^18^F]SynVesT-1, [^18^F]SynVesT-2, [^18^F]UCB-J and [^18^F]-SDM-16^3, 5^. UCB-J, -A and -H are derivatives of the anti-epileptic drug levetiracetam, known to specifically target SV2A^6, 7^. Of these compounds, [^11^C]UCB-J has shown the most promise for *in vivo* application due to its excellent grey matter uptake, fast kinetics, whole-body distribution and good reproducibility, as was shown in humans^2, 4, 8–11^, nonhuman primates (NHPs)^12^, minipigs^13^, rats^14^ and mice^15–19^. Since its discovery, [^11^C]UCB-J has made tremendous progress in clinical use. Remarkably, UCB-J has an excellent track record in clinical trials, where it has already been employed successfully in AD (NCT03493282, NCT03577262), PD (NCT04243304)^11^, HD (NCT04701580)^20^, and major depressive disorders (NCT04091971). Additionally, [^11^C]UCB-J PET studies are active or planned in schizophrenia (NCT04038840), drug use disorders (NCT04721418, NCT03527485, NCT05465538, NCT05472818), epilepsy (NCT05450822), HIV neuropathogenesis (NCT05586581), and multiple sclerosis (NCT03134716), amongst others (cf. clinicaltrials.gov). In this context, we have previously investigated the preclinical relevance of [^11^C]UCB-J PET in mouse models of HD^19^, spinal cord injury (SCI)^16^ and obsessive control disorder (OCD)^17^.

[^11^C]UCB-J has an excellent kinetic profile and reproducibility; however, the applicability of [^11^C]UCB-J PET imaging in a (pre)clinical setting remains limited due to practical limitations. C-11-labeled compounds present significant obstacles since they require an on-site cyclotron and have a shorter application window due to the short radiochemical half-life (20.38 min)^21^. Since ^18^F has a significantly longer radiochemical half-life (109.7 min), meaning fewer practical hurdles and higher image resolution, the search for an ^18^F-labeled SV2A ligand has continued in the scientific research community^22^. Due to the excellent characteristics of [^11^C]UCB-J, further research has focused on the development of F-18-labeled UCB-J analogues as well as synthesis of [^18^F]UCB-J. [^18^F]UCB-J radiosynthesis and validation in NHPs was first reported by Li *et al*., demonstrating desirable pharmacokinetics and *in vivo* binding characteristics, very similar to those of [^11^C]UCB-J^23^. [^18^F]UCB-J appears to exhibit the same kinetics and metabolism as [^11^C]UCB-J, with the added advantage of a more practical isotope and better image resolution. However, the radiochemical yield reported by Li *et al*. was extremely low (1-2%, not decay-corrected) with moderate molar activity (A_m_ = 59.0 ± 35.7 GBq/umol), hampering practical implementation^22, 23^.

Here, we report an improved [^18^F]UCB-J radiosynthesis protocol with which significantly higher radiochemical yields can be obtained and we provide an extensive validation of [^18^F]UCB-J PET in mice. This includes an assessment of [^18^F]UCB-J stability through a metabolite study, a scan duration assay, and a levetiracetam blocking study for evaluation of [^18^F]UCB-J specificity. Lastly, we examined the reproducibility of [^18^F]UCB-J *V*_T(IDIF)_ through a test-retest analysis.

## MATERIAL AND METHODS

### Animals

A total of 30 male 9-month-old C57BL/6J mice (Strain #:029928) were obtained from Jackson Laboratories (Bar Harbour, Maine, USA)) to be used for *in vivo* µPET imaging and radiometabolite analysis in plasma. Only males were included to allow adequate comparison to prior [^11^C]UCB-J and [^18^F]SynVesT-1 studies performed by our group^15, 19,21^. All mice were portosystemic shunt-free and were single-housed in individually ventilated cages under a 12 h light/dark cycle in a temperature- and humidity-controlled environment with food and water *ad libitum*. At least one week of acclimatization was guaranteed before the start of any procedure. Animals were randomly allocated to the different experimental cohorts. All experiments were performed according to the European Committee Guidelines (decree 2010/63/CEE) and reported in compliance with the ARRIVE guidelines. Experiments were approved by the Ethical Committee for Animal Testing (ECD #2020-59) at the University of Antwerp (Belgium).

### Tracer radiosynthesis

Radiosynthesis of [^18^F]UCB-J was based on work by Li *et al*.^23^. Initially, attempts to optimize copper-catalyzed radiofluorination conditions of a trimethyltin precursor using the Cu(OTf)_2_/pyridine complex system failed to yield the desired product. As a result, we turned to an alternative radiofluorination method using the racemic 4-iodonium ylide precursor for the radiosynthesis of [^18^F]UCB-J. A TRASIS AllinOne module (TRASIS, Belgium) was used for automated radiosynthesis, semi-preparative HPLC purification and formulation of [^18^F]UCB-J. Briefly, no-carrier added aqueous [^18^F]fluoride (19.05 GBq) was produced in an Eclipse HP cyclotron (Siemens), using the ^18^O(p,n)^18^F reaction by proton bombardment of [^18^O]H_2_O (Rotem Industries). Following transfer to the hot cell, the activity was trapped on a QMA cartridge (preconditioned with 5 mL 0.5M K_2_CO_3_ and 2x 5 mL ultrapure water, dried with nitrogen). [^18^F]F^−^ was eluted from the QMA cartridge to reaction vial 1 with 0.8 mL of a 7.8 mM TEAB solution in MeCN/H_2_O (90:10 v/v) and evaporated to complete dryness. After cooling to 50°C, 0.5 mL DMF and 2.5 mg 4-iodonium ylide UCB-J precursor in 0.5mL DMF were sequentially added to the reaction vial, and the reaction mixture was heated to 140°C for 10 min. Next, the crude reaction mixture was cooled to 50°C and quenched with 10 mL Water For Injection (WFI), and passed through a Waters tC18 Sep-Pak cartridge. After washing with 10 mL purified water, the cartridge was eluted with EtOH (0.5 mL) to a second vial, where it was diluted with WFI (0.5 mL) and then injected onto a semi-preparative ChiralPak IB N-5 HPLC column (10 × 250 mm, 5 μm, eluting with a mobile phase of 35% CH_3_CN and 65% 20 mM ammonium bicarbonate solution (pH 8.6) at a flow rate of 5 mL/min). The early fraction containing desired radioactive product, (R)-enantiomer of 4-[^18^F]UCB-J (retention time t_R_=25.5 min) was collected, and the product was formulated with 0.9% NaCl to obtain a radiotracer solution containing <10% EtOH to a final volume of 10 mL. The product was further sterile-filtered (0.22 µm Cathivex GV 25 mm filter unit; Merck, Belgium) to a final product vial. No free radioactive fluoride was detected in our final synthesis product. The total synthesis time was 90 min including purification and formulation. [^18^F]UCB-J was prepared with >99% radiochemical purity and a radiochemical yield of 8.5±3.2% (decay-corrected); 4.7±1.8% (not decay-corrected). The molar activity (A_m_) was 60.59±10.59 GBq/µmol (*n* = 8).

### Metabolite assessment and correction

Assessment of [^18^F]UCB-J and its radiometabolites in plasma was performed in C57BL/6J mice (*n* = 15; 2-3 per timepoint). Mice were injected intravenously (i.v.) with 5.18±0.28 MBq in 200 μl. As a terminal procedure under anesthesia (mixture of isoflurane with oxygen (5%)), blood (>300 µL)) was withdrawn via cardiac puncture at different timepoints post-injection (5, 15, 30 and 60 min).

The procedure for metabolite analysis was performed adapting a previously described radiometabolite protocol^15^ to [^18^F]UCB-J. Briefly, plasma samples (150 µL) were prepared by centrifugation of blood at 4500 × rpm (2377 x rcf) for 5 min and then mixed with equal amounts of ice-cold acetonitrile. A 10 µL aliquot of cold reference standard (1 mg/mL) was added and the mixture was subsequently centrifuged at 4500 × rpm (2377 x rcf) for 5 min to precipitate denatured proteins. The supernatant and precipitate were separated and both fractions were counted in a gamma counter (Wizard 2 2480, Perkin Elmer, USA). Following this, 100 µL of plasma supernatant was loaded onto a pre-conditioned reverse-phase (RP)-HPLC system (Phenomenex Kinetex EVO C18 5 μm HPLC column (150 x 4.6 mm) + Phenomenex security guard pre-column). Elution was performed using NaOAc 0.05M pH 5.5 and acetonitrile (65:35) buffer at a flow rate of 1 mL/min. 20 fractions were collected (0.5 min/fraction) and counts per min (CPM) were obtained from a gamma counter. To determine the recovery of [^18^F]UCB-J as well as the stability of the tracer during the workup, control experiments were performed using blood spiked *in vitro* with ∼185 kBq (∼5 µCi) of radiotracer.

Using PMOD 3.6 software (Pmod Technologies, Zurich, Switzerland), a population-based metabolite correction was generated. Individual parent fractions were averaged and fitted with a sigmoid function, since this model provided the best visual fit with lowest standard error. Individual mouse image-derived input functions (IDIFs) were obtained by extracting the signal over time (time activity curve or TAC) in the left ventricle of the heart^21^. These IDIFs were corrected with the population-based metabolite correction and with the plasma-to-whole-blood ratio.

### Dynamic µPET acquisition and reconstruction

Dynamic 120 min µPET/computed tomography (CT) images were acquired on two Siemens Inveon PET-CT scanners (Siemens Preclinical Solution, Knoxville, USA). Mice were induced and maintained under anesthesia using a mixture of isoflurane with oxygen (5% and 1.5%, respectively). Physiological parameters, i.e. respiration and temperature, were continuously monitored using a Monitoring Acquisition Module (Minerve, France) throughout preparation steps and experiments. The radiotracer was administered to the mice i.v. into the tail vein. An automated pump (Pump 11 Elite, Harvard Apparatus, USA) was employed to ensure gradual injection of the bolus (1 mL/min over a 12s interval). On average, mice were injected with 7.54 ± 2.95 MBq, corresponding to a mass of 1.97 ± 0.08 µg/kg (baseline cohort). Animal and dose parameters for baseline scans are reported in **Table S1A**. Following each two-hour µPET scan, a 10 min 80 kV/500 μA CT scan was performed for attenuation correction and for anatomical coregistration in further processing steps.

Acquired PET data was reconstructed into 45 frames of increasing length (12×10s, 3×20s, 3×30s, 3×60s, 3×150s, and 21×300s). List-mode iterative reconstruction was performed with proprietary spatially variant resolution modelling in 8 iterations and 16 subsets of the 3D ordered subset expectation maximization (OSEM 3D) algorithm^24^. Dead time correction and CT-based attenuation correction was performed. PET image frames were reconstructed on a 128×128×159 grid with 0.776×0.776×0.796 mm^3^ voxels.

### Levetiracetam blocking study

To validate *in vivo* specificity of [^18^F]UCB-J towards SV2A, mice were injected i.v. with levetiracetam (LEV)(Merck, Germany) 30 min prior to radiotracer injection as previously described^15, 21^. The high-affinity SV2A ligand LEV was administered at two different doses: 50 and 200 mg/kg (*n* = 5 and 4, respectively). Following LEV administration, 120 min dynamic µPET acquisitions were performed as described above. For each blocked scan, a baseline scan (*n* = 9) was obtained from the same animal for adequate comparison. We aimed to scan the mice twice with 7 days in between scans. Between baseline and blocking scan, we scanned the mice with 9±1.9 days (range 7-11 days; LEV50) and 9±3.4 days (range 4-11 days; LEV200) in between. Details on animal and dose parameters for the blocking study are reported in **Table S1B**. No significant difference was observed between baseline and blocking scans in terms of injected mass, injected dose, age or body weight for the LEV 50 mg/kg cohort. For the LEV 200 mg/kg cohort, only age was significantly different between baseline and blocking scan (9 days; *p*=0.0128). However, we considered the difference sufficiently small to be negligible at such advanced age (270-279 days).

### Image processing and kinetic modelling

Processing of PET data and kinetic modelling were performed using PMOD 3.6 software. Spatial normalization of the images was performed as previously reported^15, 21, 25^. Briefly, a [^18^F]UCB-J PET template was created via averaging static baseline images (*n* = 8) that were normalized to the Waxholm Atlas space. Subsequently, individual static images (*n* = 8) were spatially normalized to the [^18^F]UCB-J template and the transformation was saved for each individual scan. Given the lack of signal in the blocking images, this approach was slightly adapted. An accurate transformation could only be obtained by generating a static image out of 6 dynamic frames (from 30-80 seconds), providing adequate signal for spatial normalization. All dynamic image frames were transformed according to their individual static image transformations, and visually checked for any errors. Volumes-of-interest (VOIs) based on the Waxholm Atlas space were overlaid onto the dynamic images to extract regional TACs, specifically for the striatum (STR), motor cortex (MC), thalamus (THAL), hippocampus (HC) and the cerebellum (CB). No radioactive uptake was observed in bone (due to possible defluorination of [^18^F]UCB-J) or harderian glands that could interfere with image analysis. Regional TACs were fitted using compartmental models (one-, two- or three-tissue compartmental model (1TCM, 2TCM or 3TCM)) and the Logan plot method. For 2TCM and 3TCM, the blood volume fraction (*V*_B_) was fixed at 3.6%. For Logan, the linear phase (t*) was determined based on the model fit with a 10% maximal error with t* ranging between 10-25 min (striatum) for a scan duration of 120 min. *V*_T(IDIF)_ estimations based on 1TCM are not reported given that the model failed to fit regional TACs. 2TCM and Logan were selected as kinetic models of choice after application of IDIF correction for metabolites and plasma-to-whole-blood ratio. 2TCM allowed estimation of the microparameters *K*_1_, *k*_2_, *k*_3_ and *k*_4_ in addition to the *V*_T(IDIF)_. *K*_1_ (mL/cm^3^ per min) and *k*_2_ (1/min) represent tracer transport from plasma to tissue and back, whereas *k*_3_ (1/min) and *k*_4_ (1/min) represent tracer transport from the non-displaceable compartment to the specific compartment and back.

Parametric maps (Logan plot, *V*_T(IDIF)_) were generated with the voxel-based graphical analysis tool PXMOD (PMOD v3.6) based on the plasma activity (metabolite corrected IDIF) as input function. Individual parametric maps were averaged and overlaid onto a 3D MRI mouse brain template for anatomical reference.

### *V*_T(IDIF)_ time stability and reproducibility

Time stability of the *V*_T(IDIF)_ was evaluated for both 2TCM and Logan plot kinetic models by repeatedly excluding the last 10 min of PET acquisition from 120 min. In total, 10 scan duration outcomes were analyzed (120 min down to 30 min). The reproducibility of the *V*_T(IDIF)_ was assessed through test-retest analysis. Following an initial [^18^F]UCB-J PET scan, mice were resubjected to [^18^F]UCB-J PET after 5.4 ± 0.97 days under nearly identical conditions. Animal and dose parameters for the test-retest study are reported in **Table S1C**. No significant difference was found between T and RT in terms of injected activity (T: 8.3±3.5 MBq vs. RT: 9.8±4.1 MBq). At RT, mice had a significantly lower body weight (T: 35.5 ±2.4 g vs. RT: 34.0±2.3 g). While a significant difference was detected for injected mass (T: 1.98±0.04 µg/kg vs. RT: 2.03±0.04 µg/kg), it was sufficiently small (<5% of average injected mass) to be deemed negligible.

### Statistical analysis

Agreement between two variables, such as *V*_T(IDIF)_ obtained from different kinetic models or from different scan durations, was assessed using Pearson’s correlation and linear regression. The difference in *V*_T(IDIF)_ between baseline and blocking scans was determined by applying 2-way ANOVA with *post hoc* Bonferroni correction. Lassen plots were generated to estimate target occupancy. Test-retest analysis included the generation of a Bland-Altman plot for which the % difference was calculated as follows:

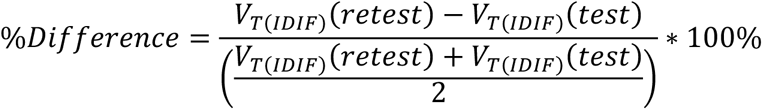

Reproducibility of the *V*_T(IDIF)_ was evaluated through the intraclass correlation coefficient (ICC) and relative and absolute test-retest variability (TRV and aTRV, respectively). The coefficient of variability (%COV = standard deviation/mean*100%) was calculated to evaluate inter-subject variability. The ICC was calculated in JMP Pro 16 (SAS Institute Inc., USA) using a mixed-model reliability analysis for absolute agreement. The (a)TRV were calculated as follows:

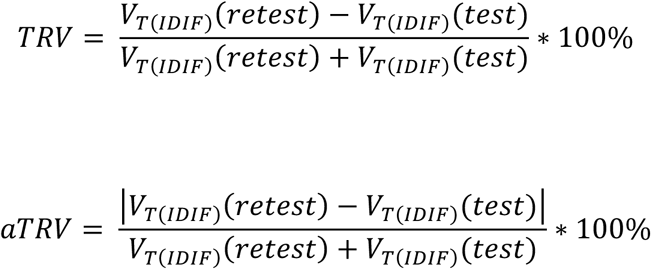

All statistical analyses were performed with GraphPad Prism (Version 9.4.1) software. All tests were two-tailed and significance was set at *p*<0.05. All data are represented as mean ± standard deviation (SD) unless otherwise specified.

## RESULTS

### Plasma radiometabolite analysis

[^18^F]UCB-J was quickly metabolized after injection in mice (*n* = 2-3 per timepoint), as depicted in **Figure 1A**. HPLC analysis of the plasma revealed at least one radiometabolite ([^18^F]M, t_R_ = 3 min) in addition to the intact radiotracer peak ([^18^F]UCB-J, t_R_ = 7 min). The percentage intact [^18^F]UCB-J was plotted over time and could be fitted with a sigmoid function, as shown in **Figure 1B**. Intact [^18^F]UCB-J accounted for 65.5 ± 3.2% at 5 min p.i., for 21.0 ± 2.7% at 15 min p.i., 12.6 ± 2.7% at 30 min p.i. and 9.6 ± 2.6% at 60 min p.i.. The plasma-to-whole blood (WB) ratio was stable over time with an average 0.88 ± 0.06% across all timepoints (**Figure S1**).

**Figure 1:**
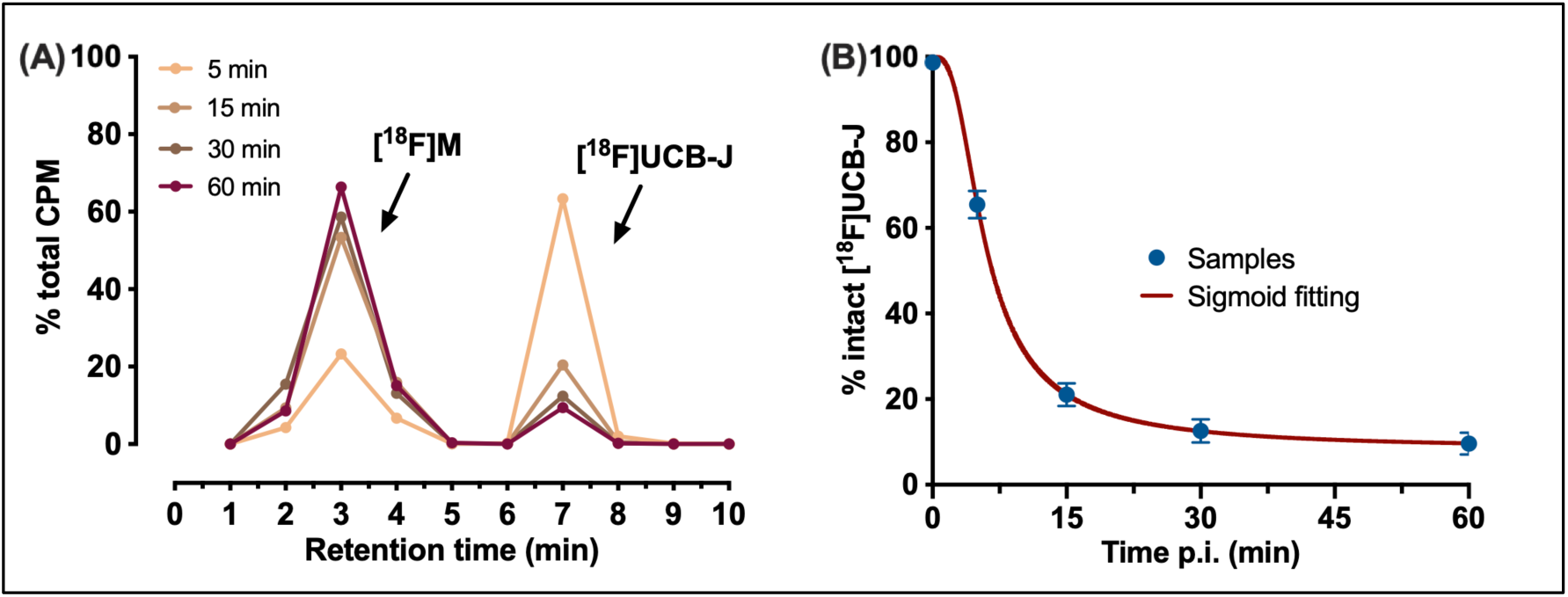
(A) Plasma elution profile at 5, 15, 30 and 60 min post [^18^F]UCB-J injection and (B) percentage intact [^18^F]UCB-J fitted with a sigmoid function. Data are presented as mean % total CPM (counts per min) of total eluted radioactivity (*n* = 2-3 per timepoint).

### Kinetic modelling

Individual IDIFs were obtained and corrected for metabolites, including a correction for the percentage intact tracer available, as well as a correction for the plasma-to-whole blood ratio (population-based)(**Figure S2A**). [^18^F]UCB-J standardized uptake value (SUV) TACs were additionally obtained from a 120 min dynamic PET acquisition for the following brain regions: STR, MC, HC, THAL and CB (**Figure S2B**). In these regions, the kinetic profile of [^18^F]UCB-J uptake indicates peak uptake around the first 10 min and reversible kinetics given the stable washout over time.

Kinetic modelling was performed with 2TCM and 3TCM using SUV TACs and the plasma metabolite-corrected IDIFs. Metabolite correction was required to achieve acceptable fitting. Both 2TCM and 3TCM provided a good visual fit, as shown in **Figure S3A and S3B**, respectively. In terms of goodness-of-fit, 3TCM provided a better Akaike Information Criterion (AIC) value than 2TCM (**Table S2**). However, given the excellent correlation between *V*_T(IDIF)_ obtained from these models (r^2^ = 0.994, *p*<0.0001)(**Figure S3C**) and that the first-order rate constants *K_1_* (representing influx from blood to tissue) were more stable with 2TCM (STR: 1.17±0.25 mL/cm^3^ per min with standard error (SE) 3.1±2.3%) compared to 3TCM (STR: 1.92±0.79 mL/cm^3^ per min with SE 14.4±10.5%), 2TCM was deemed the model of choice for [^18^F]UCB-J. Additionally, Logan graphical analysis may be employed for simplification where estimation of the microparameters is not required. Logan offers a slight underestimation of the *V*_T(IDIF)_ compared to 2TCM *V*_T(IDIF)_ but otherwise reveals an excellent correlation (r^2^ = 0.989, *p*<0.0001)(**Figure S3D**).

### [^18^F]UCB-J baseline quantification

Regional *V*_T(IDIF)_ baseline values obtained from 120 min scan duration are reported in **Table 1**. Based on 2TCM, *V*_T(IDIF)_ quantification ranged from 15.3 ± 1.8 mL/cm^3^ in the CB to 24.8 ± 3.0 mL/cm^3^ in the THAL. A comparable standard deviation was seen in all regions. Estimated *K_1_* values ranged from 1.17 ± 0.25 in the STR to 1.61 ± 0.40 in the THAL. Low standard errors were seen for *K_1_* in all regions (2.4 to 3.9%). *k*_2_ values ranged from 0.28 ± 0.14 (CB) to 0.68 ± 1.35 (MC), whereas *k*_3_ and k_4_ ranged from 0.20 ± 0.15 (CB) to 0.47 ± 0.60 (MC) and 0.060 ± 0.009 (STR) to 0.064 ± 0.014 (CB), respectively. Standard errors were relatively low for k_2_, k_3_ and k_4_ (<15%, <17%, and <8%, respectively). An overview of the 2TCM microparameters as well as their respective standard errors is reported in **Table 1**.

**Table 1:**
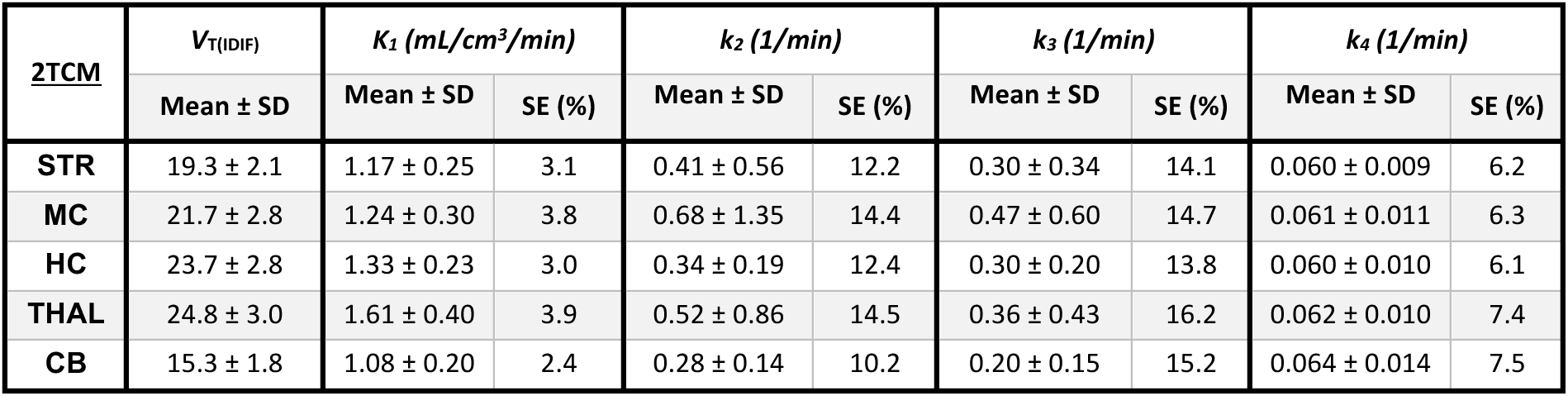
Volume of distribution (*V*_T(IDIF)_) and microparameters K_1_, k_2_, k_3_, and k_4_ based on the two-tissue compartmental model (2TCM) for the striatum (STR), motor cortex (MC), hippocampus (HC), thalamus (THAL) and cerebellum (CB). Data represented as mean ± standard deviation (SD) and standard error (SE).

### [^18^F]UCB-J time stability

To determine the minimal scan duration for accurate *V*_T(IDIF)_ quantification, the scan duration was repeatedly shortened from 120 min to 30 min (**Figure 2A**). A scan duration of 60 min was deemed appropriate given that the *V*_T(IDIF)_(Logan) change did not exceed the predetermined limit (<10% deviation per animal) while an inter-individual standard deviation of <5% was maintained. A Pearson’s correlation comparing the *V*_T(IDIF)_(Logan) output between 60 min and 120 min scans revealed significant correlation (*p*<0.0001, r^2^=0.987)(**Figure 2B**). Further shortening the scan duration to 50 min or less is not advised, given that considerable bias (>10% deviation) was seen in 3/22 mice.

**Figure 2:**
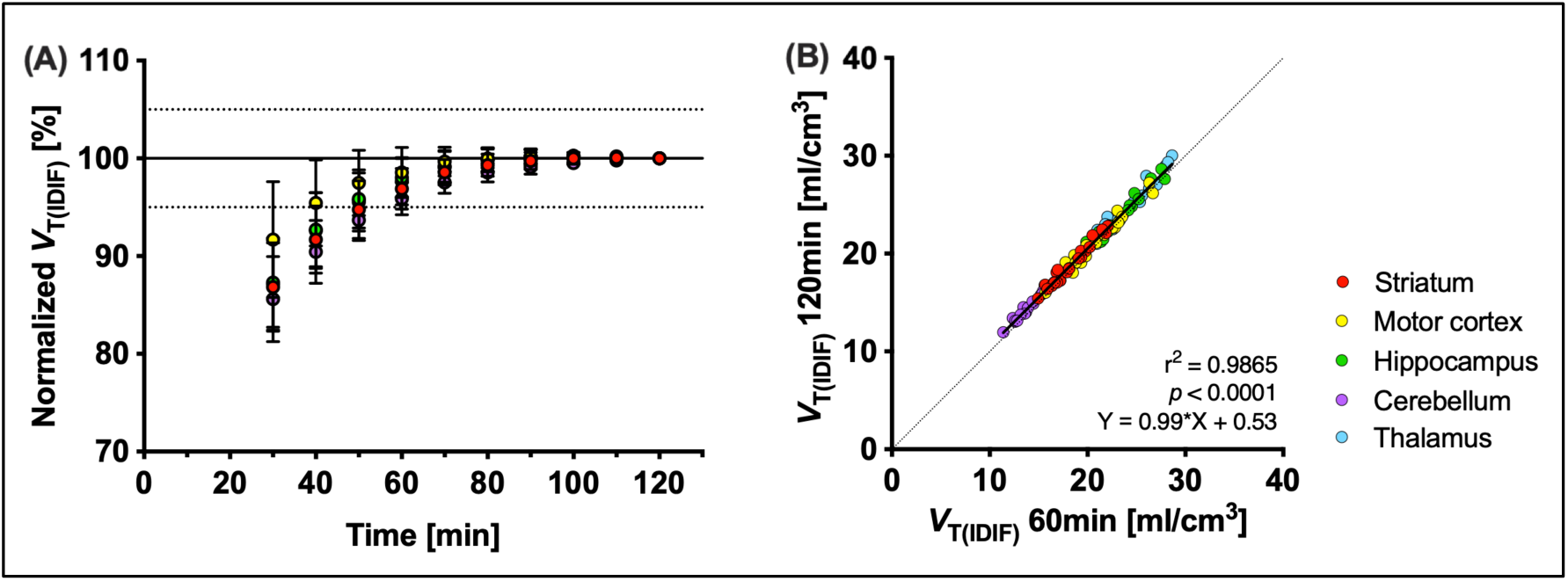
Time stability of the averaged volume of distribution metric (*V*_T(IDIF)_(Logan)) in function of scan acquisition time. (A) Time stability of the *V*_T(IDIF)_ at different acquisition times (120 to 30 min in steps of 10 min). (B) Pearson’s correlation between *V*_T(IDIF)_ calculated based on 120 min scans and *V*_T(IDIF)_ calculated based on 60 min scans. Five different brain regions were assessed: striatum, motor cortex, hippocampus, cerebellum, and the thalamus. *n* = 22.

### Levetiracetam blocking study

To evaluate the specificity of [^18^F]UCB-J binding to SV2A in mice, mice were pretreated with LEV 30 min prior to radiotracer injection. LEV pretreatment resulted in an early and steep SUV TAC decline following the initial uptake peak (**Figure 3**), indicating reduced target binding.

**Figure 3:**
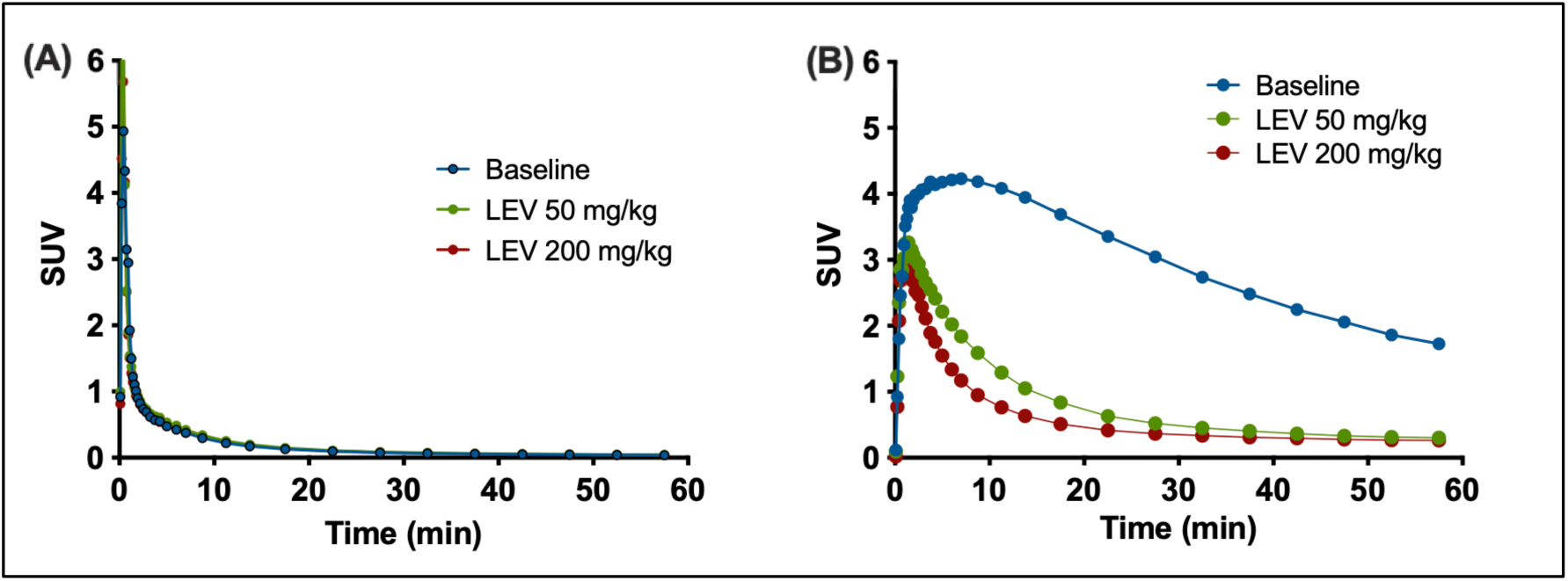
(A) Image-derived input function (IDIF) and (B) standardized uptake values time activity curves (SUV TACs) of the striatum for baseline (*n* = 9) and blocking scans (LEV 50: *n* = 5; LEV 200: *n* = 4). SUV TACs are influenced by levetiracetam (LEV) pretreatment, whereas the IDIF remained unchanged.

Baseline and blocking parametric maps based on *V*_T(IDIF)_ (Logan) are shown in **Figure 4A**, and regional *V*_T(IDIF)_ (Logan) quantification is plotted in **Figure 4B**. For a scan duration of 60 min, LEV pretreatment resulted in a dose-dependent change in *V*_T(IDIF)_ in all regions investigated. Overall, a dose of 50 mg/kg LEV resulted in a *V*_T(IDIF)_ decrease of 73.81– 82.64% over all regions investigated, whereas a dose of 200 mg/kg resulted in a decrease of 74.69–85.10% (significantly different from baseline, *p*<0.0001). Lassen plots for each dose revealed estimated SV2A occupancies of ∼79% (slope: 0.7895 at 50 mg/kg LEV) or ∼100% (slope: 1.031 at 200 mg/kg LEV) for the STR (**Figure 4C and 4D**), indicating high *in vivo* binding specificity of [^18^F]UCB-J to SV2A. The non-displaceable volume of distribution (*V*_ND_) as estimated by the Lassen plot is approximately 0.1 mL/cm^3^ (50 mg/kg LEV) and 3.7 mL/cm^3^ (200 mg/kg LEV). The effect of LEV pretreatment on the *V*_T(IDIF)_ and associated occupancy estimates for all investigated brain regions are reported in **Table 2**.

**Figure 4:**
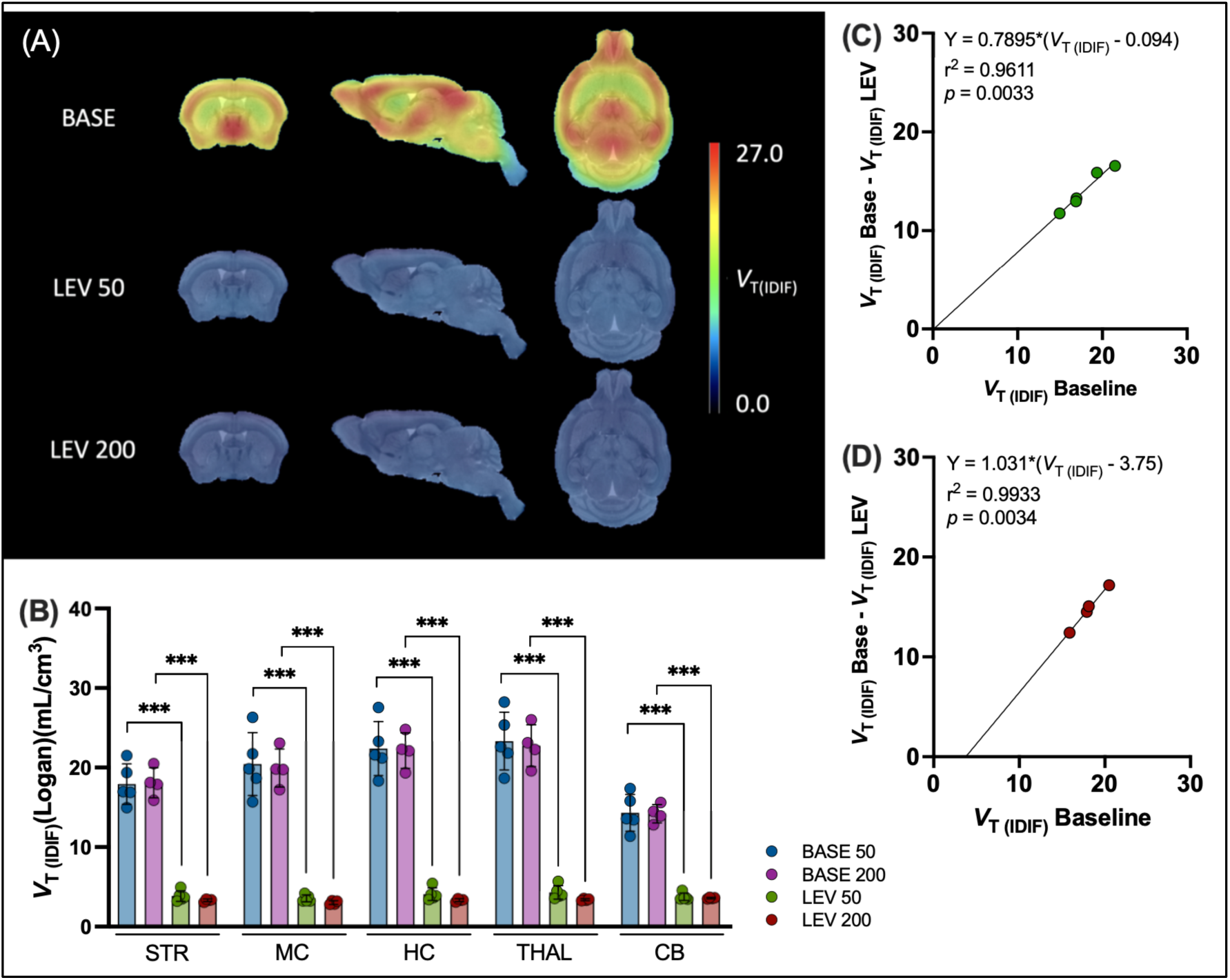
Levetiracetam (LEV) pretreatment results in a dose-dependent decrease of [^18^F]UCB-J binding. (A) Parametric maps overlaid on an MRI template for baseline, LEV 50 mg/kg and LEV 200 mg/kg blocking scans. (B) V_T(IDIF)_(Logan) quantification for baseline and blocking cohorts. A dose-dependent decrease in V_T(IDIF)_(Logan) can be noted. (C, D) Estimation of SV2A occupancy in the striatum through Lassen plots. Blocking with 50 (C) or 200 (D) mg/kg leads to an estimated SV2A occupancy of ∼79% and ∼100%, respectively. Baseline: *n* = 5 (BASE 50) and 4 (BASE 200), LEV 50: *n* = 5, LEV 200: *n* = 4.

**Table 2:** Volume of distribution (*V*_T(IDIF)_) for the striatum (STR), motor cortex (MC), hippocampus (HC), thalamus (THAL) and cerebellum (CB) based on Logan graphical analysis. Estimated occupancy based on regional Lassen plots. BASE = baseline scans (60 min), LEV = levetiracetam pretreatment (50 or 200 mg/kg). *n* = 5 (LEV 50) and 4 (LEV 200).

| **[<sup>18</sup>F]UCB-J** | **BASE LEV 50** | **LEV 50** |  | **BASE LEV 200** | **LEV 200** |  |
| --- | --- | --- | --- | --- | --- | --- |
|  | **$V_{T(IDIF)}(\text{Logan})$**<br>**Mean ± SD** | **$V_{T(IDIF)}(\text{Logan})$**<br>**Mean ± SD** | **Estimated**<br>**occupancy** | **$V_{T(IDIF)}(\text{Logan})$**<br>**Mean ± SD** | **$V_{T(IDIF)}(\text{Logan})$**<br>**Mean ± SD** | **Estimated**<br>**occupancy** |
| **STR** | 17.9 ± 2.5 | 3.9 ± 0.7 | 78.9% | 18.1 ± 1.9 | 3.3 ± 0.2 | 103.1% |
| **MC** | 20.4 ± 4.0 | 3.6 ± 0.4 | 91.8% | 20.0 ± 2.4 | 3.0 ± 0.2 | 98.2% |
| **HC** | 22.4 ± 3.4 | 4.1 ± 0.8 | 79.1% | 22.1 ± 2.2 | 3.3 ± 0.2 | 104.0% |
| **THAL** | 23.3 ± 3.6 | 4.3 ± 0.9 | 81.4% | 22.8 ± 2.6 | 3.4 ± 0.1 | 98.5% |
| **CB** | 14.3 ± 2.3 | 3.8 ± 0.5 | 86.8% | 14.2 ± 1.2 | 3.6 ± 0.1 | 103.8% |

### [^18^F]UCB-J test-retest

Reliability of [^18^F]UCB-J *V*_T(IDIF)_ (Logan, 60 min) was assessed through a test-retest (T-RT) analysis. Regional *V*_T(IDIF)_(Logan) values did not significantly differ between T and RT based on 2-way ANOVA with *post-hoc* Bonferroni correction. Parametric maps show the same distribution for T and RT (**Figure 5A**). A Bland-Altman analysis suggested a bias of 9.12 and 95% limits of agreement from −14.25% to 32.48% with no region-specific trend (**Figure 5C**). Accordingly, relative TRVs are positive and stable over the STR, MC, HC, THAL, and CB. The highest absolute TRVs were found in the MC (6.2±5.0%) and THAL (6.1±5.2%). Regional ICC values ranged between 0.66 (THAL) and 0.75 (STR), indicating ‘moderate agreement’ (**Table S3**)^26^. This agreement was confirmed by a Pearson’s correlation assay, in which T and RT *V*_T(IDIF)_(Logan) values showed significant correlation (r^2^ = 0.773, *p*<0.0001)(**Figure 5B**).

**Figure 5:**
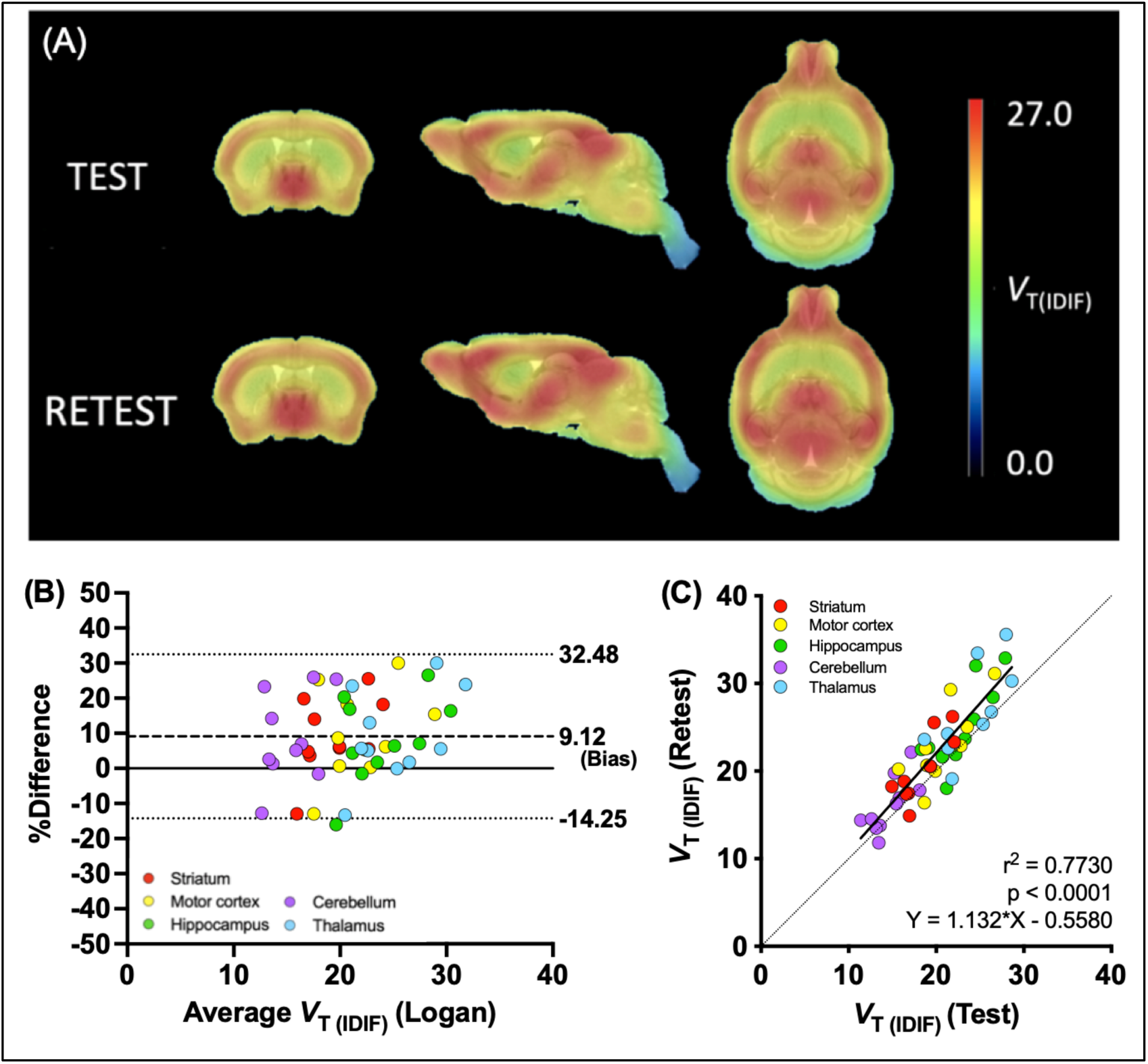
(A) Parametric maps showing a similar distribution of radiotracer in test and retest scans. (B) Bland-Altman plot reports a bias of 9.12 and 95% limits of agreement from −14.25% to 32.48%. (C) Pearson’s correlation between the *V*_T(IDIF)_(Logan) from test and retest scans within the same mice. (*n* = 10 mice; 5 regions).

## DISCUSSION

Given the biological relevance of synaptic communication in a wide variety of diseases, non-invasive monitoring of SV2A as a proxy for synaptic density quantification is vital to better understand disease pathology and to monitor disease progression and therapeutic efficacy. Several SV2A PET tracers have been developed for this purpose, of which [^11^C]UCB-J is currently the most used radiotracer. We previously validated the use of [^11^C]UCB-J in mice, demonstrating favorable metabolism, selectivity to SV2A and optimal kinetics^15^. Although these *in vivo* properties are excellent, practical [^11^C]UCB-J radiosynthesis remains challenging because of the short radiochemical half-life of its ^11^C isotope. With the aim of preserving the excellent properties of UCB-J and gaining the advantage of a longer-lived isotope, we set out to improve the production protocol for [^18^F]UCB-J and to validate [^18^F]UCB-J in a preclinical context. We established a fully automated cassette-based radiosynthesis process that can be easily adopted for the clinical translation and in the implementation of [^18^F]UCB-J in other research facilities. We optimized the radiofluorination procedure by Li *et al.* (2019)^23^ by decreasing the amount of base (TEAB) and the reaction temperature to obtain the best incorporation of ^18^F-fluoride and reduced racemization, leading to improved radiochemical yields. The A_m_ reported by Li *et al*. was moderate (59.0±35.7 GBq/µmol)^23^. For current study, we obtained similar but more stable A_m_ (60.59±10.59 GBq/µmol), which was sufficient for satisfactory image quality and multiple scan runs. For [^11^C]UCB-J, we previously reported molar activities of 96.5±13.3 GBq/µmol^19^ and 78.8±23.0 GBq/µmol^15^, both of which are in line with what we here achieved for [^18^F]UCB-J. Notably, although our A_m_ is generally lower for [^18^F]UCB-J compared to [^11^C]UCB-J, the ^18^F isotope has the great advantage of slower decay, entailing fewer productions with multiple runs.

Structurally, [^18^F]UCB-J and [^11^C]UCB-J are identical molecules, the only difference being the isotope used for radiolabeling^5^. Given their equivalent structure and *in vivo* behavior in NHPs, similar physiobiological *in vivo* behavior of [^18^F]UCB-J and [^11^C]UCB-J was anticipated in mice^23^. Plasma analysis of [^18^F]UCB-J revealed at least one polar metabolite and rapid plasma metabolism, as observed for [^11^C]UCB-J^15^. The percentage intact tracer over time in plasma for [^18^F]UCB-J was also in agreement with what we previously found in mice: 61.1±6.8% ([^11^C]UCB-J: 65.5±3.2%), 22.5±4.2% (21.0±2.7%), 13.6±3.7% (12.6±2.7%) at 5, 15 and 30 min p.i., respectively^15^. Since these characteristics are similar, we may expect the kinetic profile and metabolism of [^18^F]UCB-J in humans to be comparable to that of [^11^C]UCB-J, which has already shown to be exemplary^2, 27^.

[^18^F]UCB-J kinetics could be described by multiple kinetic models based on an IDIF, including 3TCM, 2TCM and Logan. Microparameter (*K*_1_, *k_2_*, *k*_3_ and *k*_4_) estimates were most stable with 2TCM. Kinetic modelling of [^18^F]UCB-J could not be performed with 1TCM, with or without metabolite correction of the IDIF. Conversely, 1TCM and Logan (without IDIF correction) were presented as models of choice for [^11^C]UCB-J modeling in the same mouse strain of similar age^15^. Notably, in current [^18^F]UCB-J study, the IDIF approach was not validated against an arterial input function (AIF). However, we previously reported such validation for [^18^F]SynVesT-1, a structurally similar radiotracer which exhibits comparable kinetics in mice^21, 28^. Accordingly, we did not encounter challenges using an IDIF-based *V*_T(IDIF)_ in terms of pharmacokinetic model fit or bias due to myocardium uptake. Importantly, the IDIF remains an estimation of the AIF. In order to compare between studies, the same method should be applied (e.g. similarly (un)corrected IDIF).

We obtained reliable *V*_T(IDIF)_ values for all relevant brain regions (STR, MC, HC, THAL and CB). Shortening the scan duration to 60 min did not significantly impact the *V*_T(IDIF),_ as evidenced by the near-perfect correlation between *V*_T(IDIF)_(120 min) and *V*_T(IDIF)_(60 min) (i.e. Logan: r^2^=0.987). These results were consistent with prior findings from our lab for other SV2A tracers, recommending 60 min scans for [^11^C]UCB-J and [^18^F]SynVesT-1^15, 21^. The *V*_T(IDIF)_ represents both the specific volume of distribution (*V*_S_) and the non-displaceable volume of distribution (*V*_ND_). We conducted a blocking study with LEV (50 or 200 mg/kg) to determine the contribution of the non-displaceable component (*V*_ND_) and thereby evaluate the affinity of [^18^F]UCB-J to SV2A *in vivo*. Our results confirmed [^18^F]UCB-J specificity to SV2A throughout the grey matter by the ubiquitous and dose-dependent decrease in *V*_T(IDIF)_(60 min) following LEV pretreatment. In the STR, SV2A occupancy was estimated at ∼79% and ∼100% for 50 and 200 mg/kg LEV, respectively, resembling that of [^11^C]UCB-J (79.2% and 97.3%)^15^. The SV2A affinity of [^11^C]UCB-J in mice appears similar despite the change in isotope. Interestingly, in NHPs, SV2A occupancies of 79% ([^18^F]UCB-J) and 87% ([^11^C]UCB-J) were reported using the same concentration of blocking agent (0.15 mg/kg UCB-J)^12, 23^. Our [^18^F]UCB-J *V*_ND_ estimations in the STR mice were low (0.1 mL/cm^3^ and 3.7 mL/cm^3^ for 50 and 200 mg/kg LEV, respectively) like those previously reported ([^11^C]UCB-J: 0.17 and 0.60 mL/cm^3^)^15, 21^.

Although the small sample size in this component of our study may confound accurate estimation of *V*_ND_, we can conclude that the contribution of *V*_ND_ to the *V*_T(IDIF)_ is limited. We further assessed [^18^F]UCB-J in terms of reproducibility, i.e. robustness of the outcome parameter *V*_T(IDIF)_. Between T and RT, [^18^F]UCB-J *V*_T(IDIF)_ values were consistent with low TRV (4.1-4.9%) and low aTRV (5.8-6.2%). The observed extent of variability was in line with the average regional aTRV of 5.3±4.1% for [^11^C]UCB-J in humans as reported by Finnema *et al*^8^. In addition to the low variability, we observed no significant difference between T and RT *V*_T(IDIF)_ values. *V*_T(IDIF)_ values significantly correlated between T and RT and were classified as moderately agreeing based on the ICC (0.66-0.75)^26^. Notably, given that the standard deviation on the regional *V*_T(IDIF)_ averages increases with decreasing scan duration (cf. [^18^F]UCB-J time stability), a smaller bias and improved correlation can be expected with longer acquisition times due to the more stable regional *V*_T(IDIF)_. Where *V*_T(IDIF)_ stability is the prime concern, scan durations from 60 min remain acceptable as evidenced by our time stability study. To our knowledge, this study is the first to report [^18^F]UCB-J T-RT properties. Overall, [^18^F]UCB-J showed acceptable reliability and reproducibility between T and RT.

## CONCLUSION

In conclusion, [^18^F]UCB-J is a selective SV2A tracer that can be modelled with 2TCM or Logan plot in mice. SV2A density could reliably and reproducibly be estimated in a non-invasive manner using an IDIF approach. [^18^F]UCB-J exhibits the same attractive properties as [^11^C]UCB-J, with the added benefit of being radiolabelled with a longer-lived isotope. Further investigation is required to explore the potential of [^18^F]UCB-J PET within neuropsychiatric and neurodegenerative disorders such as HD and schizophrenia, pathologies with known SV2A alterations.

## ACKNOWLEDGEMENTS

The authors thank Ming Min Hsia and John Mangette (Curia Global) for the synthesis of the iodonium ylide [^18^F]-precursor; Philippe Joye, Caroline Berghmans, Eleni Van der Hallen, and Annemie Van Eetveldt of the Molecular Imaging Center Antwerp (MICA) for their valuable assistance. The authors also thank Brenda Lager (CHDI) for animal supply, Maya Bader and Braden Boone (CHDI) for program management, and Simon Noble (CHDI) for reviewing the manuscript. Daniele Bertoglio, Steven Staelens and Liesbeth Everix are members of the µNeuro Research Centre of Excellence at the University of Antwerp. The University of Antwerp also funded the work through an assistant professor position for FE and DB and a full professor position for SS.

## AUTHORS’ CONTRIBUTIONS

Study conceptualization: LE, FE, LL, JB, SS, and DB

Data acquisition: LE

Methodology: LE, FE, AM, SS, DB

Data analysis and visualization: LE, AM

Result interpretation: LE, FE, LL, JB, SS, DB

Writing original draft: LE

Review and editing: LE, FE, AM, VK, LL, JB, SS, and DB

Final approval: All authors

## COMPETING INTERESTS STATEMENT

This work was funded by CHDI Foundation, Inc., a privately-funded nonprofit biomedical research organization exclusively dedicated to collaboratively developing therapeutics that improve the lives of those affected by Huntington’s disease. VK, LL, and JB are employed by CHDI Management, Inc. to manage the scientific activities of CHDI Foundation, Inc. No other potential conflicts of interest relevant to this article exist. No potential conflicts of interest with respect to the research, authorship, and/or publication of this article are present.

## MATERIAL AVAILABILITY

All data associated with this study are presented in the paper or the Supplementary Materials. Any request for material reported in this study will be available through a material and/or data transfer agreement.

## SUPPLEMENTARY MATERIAL

Supplemental material for this article is available online.

## Notes

### Summary of Updates

We 1) better defined the conditions of the blocking study (material and methods); 2) elaborated on the effects of the radiosynthesis optimization (discussion); and 3) evaluated potential spill-in effects from nearby anatomical regions (none observed, material and methods). Minor changes to improve readability or consistency (e.g. rounding up digits after the comma or making units consistent) have also been performed. Amended figure: 4B (BASE groups) Amended tables: 1,2 and S3 (digits) + S1 (BASE groups + digits)

