## Supplemental material for "Preclinical validation and kinetic modelling of the SV2A PET ligand [^18^F]UCB-J in mice"

Liesbeth Everix[1], Filipe Elvas[2], Alan Miranda Menchaca[1], Vinod Khetarpal[3], Longbin Liu[3], Jonathan Bard[3], Steven Staelens[1], Daniele Bertoglio[1, 4,5]

*[1]Molecular Imaging Center Antwerp (MICA), University of Antwerp, Wilrijk, Belgium*
*[2]University of Antwerp, Department of Nuclear Medicine, Wilrijk, Belgium*
*[3]CHDI Management, Inc., the company that manages the scientific activities of CHDI Foundation, Inc. Princeton, NJ, USA*
*[4]Bio-Imaging Lab, University of Antwerp, Wilrijk, Belgium*
*[5]μNeuro Center for Excellence, University of Antwerp, Antwerp, Belgium*

**Corresponding author:**
Prof. Daniele Bertoglio
Bio-Imaging Lab
University of Antwerp, Universiteitsplein 1, Wilrijk, Belgium


***__Table S1__**: Animal and dose parameters for the baseline study (1A), the blocking experiment (1B) and the test-retest experiment (1C). p-values refer to statistical difference between the current cell and the grayed-out cell above within the same category (e.g. 1A, 1B, 1C) based on paired t-tests. Data shown as mean ± standard deviation.*

| | | **Injected activity (MBq)** | **Molar activity (GBq/µmol)** | **Weight (gram)** | **Age (days)** | **Injected mass (µg/kg)** |
|---|---|---|---|---|---|---|
| **1A** | Baseline *(n = 22)* | 7.5 ± 2.9 | 58.6 ± 12.7 | 35.5 ± 3.0 | 273.4 ± 12.3 | 1.97 ± 0.08 |
| **1B** | Baseline LEV 50 *(n = 5)* | 9.2 ± 3.0 | 63.4 ± 3.0 | 35.3 ± 2.5 | 280.8 ± 18.5 | 1.97 ± 0.03 |
| | LEV 50 *(n = 5)* | 5.9 ± 1.6 (*p*=0.0679) | 60.9 ± 15.9 (*p*=0.7118) | 34.3 ± 1.9 (*p*=0.1810) | 284.2 ± 9.4 (*p*=0.4685) | 1.96 ± 0.04 (*p*=0.2030) |
| | Baseline LEV 200 *(n = 4)* | 6.2 ± 2.6 | 57.4 ± 16.6 | 34.0 ± 4.2 | 270.0 ± 7.9 | 1.99 ± 0.04 |
| | LEV 200 *(n = 4)* | 6.0 ± 1.8 (*p*=0.7474) | 57.1 ± 15.6 (*p*=0.9670) | 31.9 ± 2.7 (*p*=0.1174) | 279.0 ± 10.3 (*p*=0.0128) | 1.99 ± 0.03 (*p*=0.8001) |
| **1C** | Test *(n = 10)* | 8.3 ± 3.5 | 62.9 ± 3.5 | 35.5 ± 2.4 | 268.3 ± 7.2 | 1.98 ± 0.04 |
| | Retest *(n = 10)* | 9.8 ± 4.1 (*p*=0.1155) | 68.1 ± 5.4 (*p*<0.0001) | 34.0 ± 2.3 (*p*<0.0001) | 273.7 ± 7.7 (*p*<0.0001) | 2.03 ± 0.04 (*p*=0.0027) |

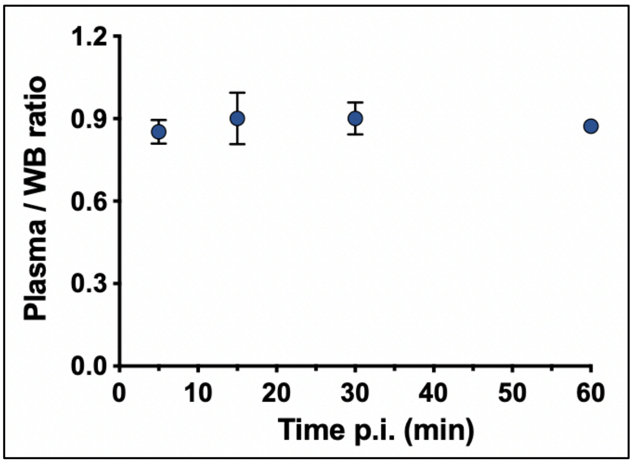


***<u>Figure S1</u>**: The plasma-to-whole blood ratio was stable around a ratio of 0.88±0.06 at all timepoints (0.85±0.04, 0.90±0.09, 0.90±0.06 and 0.87±0.01 at 5, 15, 30 and 60 min., respectively)*

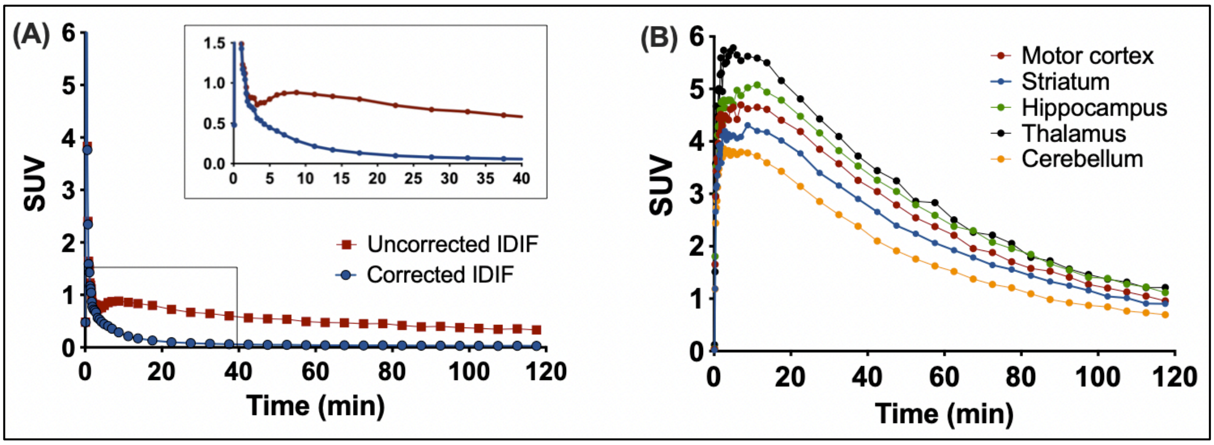


***<u>Figure S2</u>**: (A) Comparison uncorrected and corrected IDIF for [$^{18}$F]UCB-J and (B) Standardized uptake value time activity curves (SUV TACs) in five different brain regions based on an acquisition time of 120 min. n = 1.*

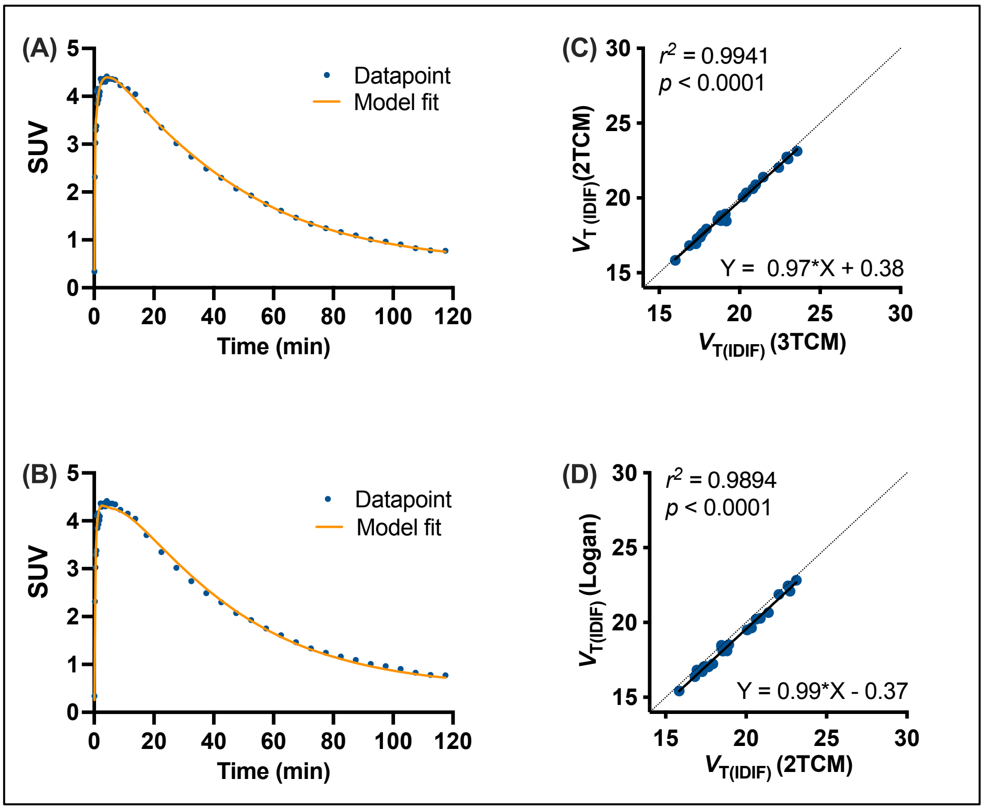


***<u>Figure S3</u>**: Representative model fits with (A) the three-tissue compartmental model (3TCM)(n = 1) and (B) the two-tissue compartmental model (2TCM)(n = 1). A Pearson's correlation presents the correlation between the $V_{T(IDIF)}$ obtained from (C) 3TCM and 2TCM (n = 22) or (D) 2TCM and Logan (n = 22). SUV = Standardized uptake value.*

***<u>Table S2</u>***: *Akaike (AIC) values for compartmental models (2TCM and 3TCM). Lower AIC values represent an improved goodness-of-fit. Data shown as mean ± standard deviation (SD).*

| **[$^{18}$F]UCB-J WT (*n* = 22)** | **AIC (2TCM) – 120 min Mean ± SD** | **AIC (3TCM) – 120 min Mean ± SD** |
|---|---|---|
| **STR** | 5.37 ± 18.73 | -17.80 ± 18.93 |
| **MC** | 2.66 ± 21.20 | -13.45 ± 24.26 |
| **HC** | 9.90 ± 16.59 | -22.79 ± 13.69 |
| **THAL** | 21.56 ± 22.40 | 2.56 ± 15.86 |
| **CB** | 10.83 ± 22.16 | -23.03 ± 15.29 |

***Table S3***: *Test and retest $V_{T(IDIF)}$(Logan) values and their associated % coefficient of variation (%COV). Regional relative and absolute test-retest variability (TRV and aTRV, respectively) values and regional intraclass correlation coefficients (ICC) are additionally reported. n = 10 mice (5 regions). CB = cerebellum, HC = hippocampus, MC = motor cortex, STR = striatum, THAL = thalamus.*

| | Test | | Retest | | | | |
|---|---|---|---|---|---|---|---|
| | $V_{T(IDIF)}$ | %COV | $V_{T(IDIF)}$ | %COV | TRV (%) | aTRV (%) | ICC |
| **STR** | 18.4 ± 2.4 | 13.2 | 20.3 ± 3.7 | 18.3 | 4.5 ± 5.4 | 5.8 ± 3.8 | 0.75 |
| **MC** | 20.8 ± 3.1 | 14.9 | 23.1 ± 4.4 | 19.1 | 4.9 ± 6.4 | 6.2 ± 5.0 | 0.72 |
| **HC** | 22.8 ± 3.1 | 13.6 | 25.0 ± 4.8 | 19.2 | 4.1 ± 6.2 | 5.9 ± 4.3 | 0.72 |
| **THAL** | 23.8 ± 3.3 | 13.7 | 26.4 ± 5.2 | 19.6 | 4.8 ± 6.6 | 6.1 ± 5.2 | 0.66 |
| **CB** | 14.6 ± 2.1 | 14.5 | 16.1 ± 3.2 | 19.6 | 4.5 ± 6.4 | 6.0 ± 5.0 | 0.69 |
